# *In vivo* widefield calcium imaging of the mouse cortex for analysis of network connectivity in health and brain disease

**DOI:** 10.1101/459941

**Authors:** Julia V Cramer, Benno Gesierich, Stefan Roth, Martin Dichgans, Marco Düring, Arthur Liesz

**Affiliations:** Institute for Stroke and Dementia Research, University Hospital, LMU Munich, Munich, Germany; Munich Cluster for Systems Neurology (SyNergy), 80336 Munich, Germany

**Keywords:** *in vivo* imaging, mouse models, neuronal network connectivity, stroke, recovery, ICA

## Abstract

The organization of brain areas in functionally connected networks, their dynamic changes, and perturbations in disease states are subject of extensive investigations. Research on functional networks in humans predominantly uses functional magnetic resonance imaging (fMRI). However, adopting fMRI and other functional imaging methods to mice, the most widely used model to study brain physiology and disease, poses major technical challenges and faces important limitations. Hence, there is great demand for alternative imaging modalities for network characterization. Here, we present a refined protocol for *in vivo* widefield calcium imaging of both cerebral hemispheres in mice expressing a calcium sensor in excitatory neurons. We implemented a stringent protocol for minimizing anesthesia and excluding movement artifacts which both imposed problems in previous approaches. We further adopted a method for unbiased identification of functional cortical areas using independent component analysis (ICA) on resting-state imaging data. Biological relevance of identified components was confirmed using stimulus-dependent cortical activation. To explore this novel approach in a model of focal brain injury, we induced photothrombotic lesions of the motor cortex, determined changes in inter- and intrahemispheric connectivity at multiple time points up to 56 days post-stroke and correlated them with behavioral deficits. We observed a severe loss in interhemispheric connectivity after stroke, which was partially restored in the chronic phase and associated with corresponding behavioral motor deficits. Taken together, we present an improved widefield calcium imaging tool accounting for anesthesia and movement artifacts, adopting an advanced analysis pipeline based on human fMRI algorithms and with superior sensitivity to recovery mechanisms in mouse models compared to behavioral tests. This tool will enable new studies on interhemispheric connectivity in murine models with comparability to human imaging studies for a wide spectrum of neuroscience applications in health and disease.

## Introduction

Functional brain networks and their alterations are of great interest for understanding mechanisms in health and neurological diseases. Connectivity analysis based on functional magnetic resonance imaging (fMRI) in humans has facilitated our growing understanding of physiological and pathological processes in the brain. However, mechanistic studies, identification of novel therapeutic targets and drug development are mostly conducted in mouse models. These models allow genetic modifications, in depth analysis of biological samples and pharmacological testing, which is driving translational sciences. To improve comparability of translational neuroscience research to findings in humans, development of brain network analysis in mice is necessary. Yet, fMRI analysis is very challenging and limited in mice, due to their size and long procedure times often combined with intravenous anesthesia and intubation (1,2). Additionally, fMRI analysis is based on blood oxygenation levels, allowing only indirect measurements of neural activity. Therefore, our aim was to establish an imaging tool that would circumvent these limitations while maintaining the comparability of the analysis approaches to human fMRI.

For this reason, we adapted *in vivo* widefield calcium imaging of neuronal activity (3,4) as a reversed translational approach from bedside to bench. We used Thy1GCaMP6s heterozygous mice, which express genetically-encoded calcium indicators (GECIs). The GECI expressed in these animals is a calmodulin-bound green fluorescent protein (GCaMP), which increases fluorescence at high intracellular calcium levels. The Thy1 promotor is limiting the GCaMP expression specifically to excitatory neurons, predominantly in layer 5 pyramidal cells (5). Hence, fluorescence intensity recorded from these mice directly mirrors cortical neuronal activity. This method not only allows repetitive, non-invasive measurements but is also blood flow-independent. This is mandatory to reduce interference of possible pathological changes in neurovascular coupling, which is understood to play a pivotal role in several brain diseases (6). Using this widefield imaging approach, we were able to repetitively acquire both spontaneous resting-state and stimulus-dependent activity from both forebrain hemispheres in healthy mice and after experimental stroke induction. We then used analysis algorithms comparable to the ones commonly used in fMRI analysis in order to generate a truly translational research tool.

Compared to previous attempts of establishing such a method, we advanced the sedation protocols to avoid anesthesia-associated bias and used objective measures to control for anesthesia artifacts. We introduced a procedure to register brains spatially between different recordings within the same mouse and between different mice and thereby ensured inter- and intra-subject comparability – a prerequisite for reliable group analysis. Furthermore, we implemented advanced analytical paradigms to neuronal calcium-imaging, such as the use of independent component analysis (ICA) for functional network characterization.

We then applied this optimized method to investigate post-stroke dynamic changes of functional connectivity as a hallmark of post-stroke brain injury and regeneration. Previous studies using fMRI in humans have advanced our knowledge about neuronal plasticity and recovery after stroke (7). Not only the changes in functional networks in the brain during acute and chronic stroke but also their influence on motoric rehabilitation have been well studied (8,9), linking functional network connectivity to stroke outcome (10–12). In our model, we specifically targeted the motor cortex in mice by using a photothrombotic stroke model and repetitively imaged animals during their recovery up to 2 months after stroke. We discovered desynchronisation of interhemispheric homotypic areas and partial restoration of network function in the chronic phases after stroke. Hence, we provide evidence for the applicability of an advanced calcium-dependent imaging modality of neuronal activity of nearly the entire forebrain cortex in mice and its feasibility for investigating the integrity and pathological/adaptive changes of functional cortical networks in acute and chronic brain diseases.

## Material and methods

### Sub-studies: the anesthesia, network and stroke study

This study consisted of three sub-studies. In a first study (further called anesthesia study), three different anesthesia protocols were tested (1.5% ISO, 1.0% ISO, MED+ISO; n=9), in order to select a protocol providing as-low-as-possible and consistent levels of anesthesia depth across animals and days. Also, the effect of each anesthesia protocol on functional connectivity was assessed.

In a second study (further called network study), ICA (in particular an extension of classical ICA called independent vector analysis (IVA)) was used to define distinct functional cortical networks on resting-state acquisitions (n=41-47). Somatosensory stimulation (n=23) was further used to identify the functional involvement of the different IVA derived networks. Regions of interest (ROI) were derived from the peak pixel of the areas being part of the independent component spatial maps.

In a third study (further called stroke study), the established anesthesia protocol and ROI were used to assess pathological changes and recovery of functional connectivity after an acute injury (i.e. photothrombosis) to the motor cortex (sham n=17; stroke n=23). This last study was meant to proof the concept that our new paradigm can be a valuable tool in the study of disease models.

### Animals

All experiments were conducted in accordance with national guidelines for the use of experimental animals and all protocols were approved by the German governmental committees (Regierung von Oberbayern, Munich, Germany). C57BL/6J-Tg(Thy1-GCaMP6s)GP4.12Dkim/J (5) heterozygous mice were bred at the Institute for Stroke and Dementia Research, Munich. Mice were housed at controlled temperature (22 ± 2 °C), with a 12-h light–dark cycle period and had access to pelleted food and water ad libitum. Both male and female animals were used for this study at 12-15 weeks of age. During all anesthetized procedures body temperature was maintained using a feedback-controlled heating system. After end of anesthesia animals were put in a heating chamber until they recovered from anesthesia.

### Skull preparation

Skull preparation was performed using a modified version of the previously published protocol of Silasi et al. (4). Briefly, after induction of anesthesia mice were placed into a stereotactic frame (product no.: 51501, Stoelting, Europe). The head was fixed, scalped and the underlying connective tissue was gently removed. After cleaning and disinfection, a layer of transparent dental cement (Quick Base S398, L-Powder clear S399, Universal Catalyst S371, Parkell C&B metabond, USA) was applied and covered with a tailored coverslip (24×60mm #1,5, Menzel-Gläser, Germany). The mice stayed anesthetized until the cement was dried and were afterwards allowed to recover for more than 48h before the first image acquisition.

### Anesthesia

In a first study three different anesthesia protocols were tested. Here, these protocols will be referred to as 1.5% ISO, 1.0% ISO and MED+ISO. The 1.5% ISO and 1.0% ISO protocols were different only in the percentage of isoflurane (1.5% and 1.0%) used to maintain different depth of anesthesia. In the MED+ISO protocol we injected 0.5mg/kg body weight of medetomidine intraperitoneally 5 minutes prior to inducing anesthesia and the isoflurane level was set at 0.75%. In all three protocols anesthesia was induced with 5% isoflurane for exactly 70 seconds, followed by 140 seconds of 1.5% ISO and finally the reduced level according to the respective protocol for at least 4 minutes for maintenance of steady-state before start of data acquisition. Isoflurane was vaporized in 30% O2 and 70% N2O. Each mouse (n=9) was imaged at least three times per condition in a randomized manner.

To define functional networks (network study) and to assess changes in post-stroke functional connectivity (stroke study) only the MED+ISO protocol was used.

### Imaging setup

*In vivo* widefield calcium imaging was performed with anesthetized animals being placed in a stereotactic frame below a customized imaging setup (**Fig. 1a**): A light beam of blue light emitting diodes (LED) of 450 nm wavelength (SOLIS 445B/M, Thorlabs, USA) and constant 650 mA current (Advanced Solis LED Driver, Thorlabs, USA) was redirected by a 495 nm dichroic longpass filter (Optical Imaging Ltd, Israel) into the optical path in front of the camera and through the chronic window on top of the skull into the cortex. The emitted light from the cortical GCaMP6s proteins passed through a set of two video lenses (NIKKOR, 85 mm f1.4, and NIKKOR 50 mm f1.2; Nikon, Japan) and a 495 nm dichroic longpass filter and a 515 nm longpass filter (both Optical Imaging Ltd, Israel). Working distance between the mouse cortex and first video lense was approximately 4.4cm. The projected image was recorded by a 2/3″ Interline CCD camera with 7.4 ×7.4 μm pixel size (Adimec 1000-m/D, Adimec, Netherlands) and using a customized longDaq software (Optical Imaging Ltd, Israel) (**Fig. 1b**). The camera field-of-view covered a quadratic area of approximately 12×12 mm = 144 mm^2^. Data was spatially binned at 3×3 pixels, resulting in an image matrix of 330×330 pixels (**Fig.1c**).

**Figure 1:**
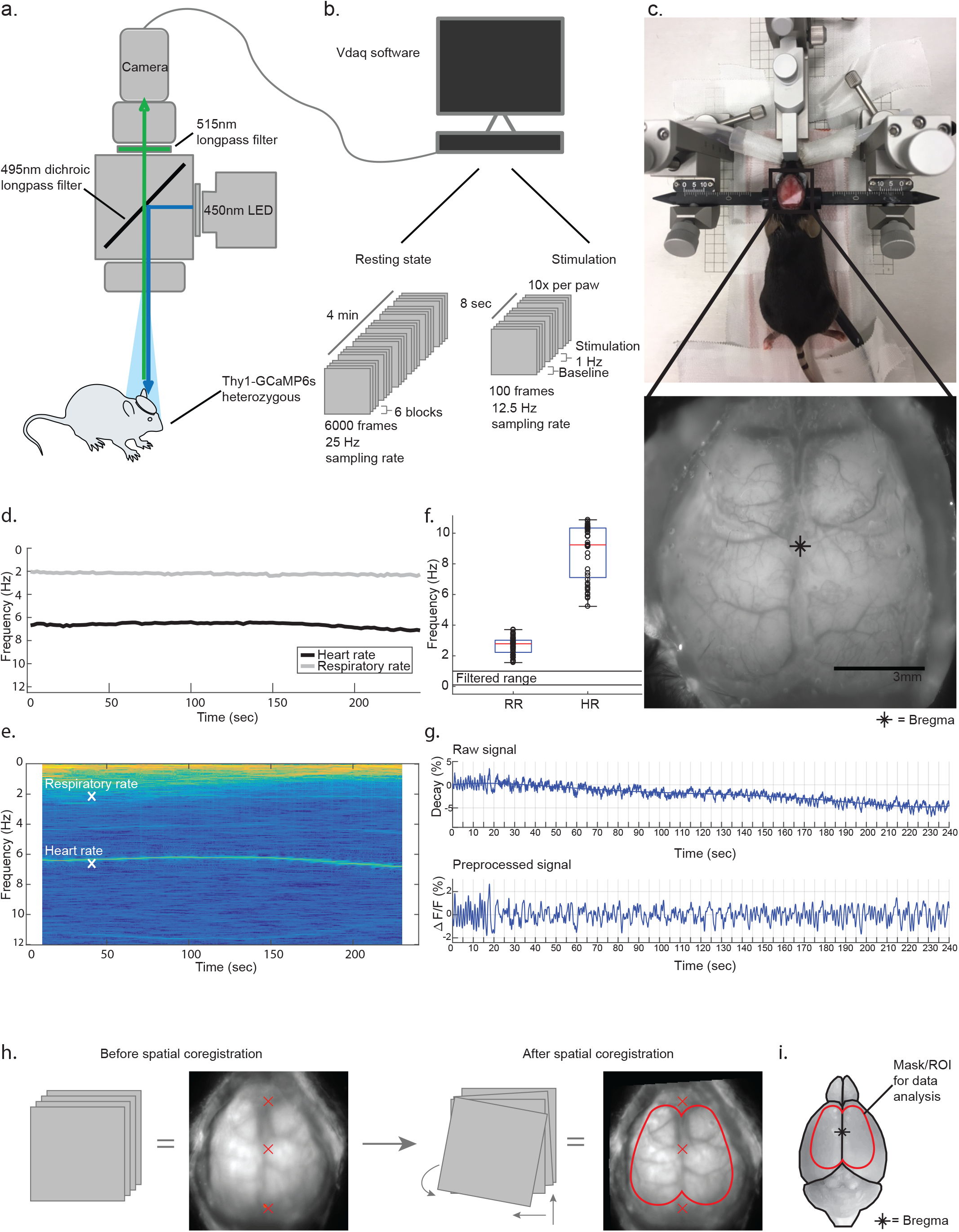
Imaging setup, data acquisition and processing steps. **a.** Customized imaging setup built with 450 nm LED light source which was directed via 495nm dichroic longpass filter to illuminate the mouse cortex; emitted GFP light was filtered by a 515 nm longpass filter before being acquired by a CCD camera. **b.** Resting-state data was recorded in 6 blocks à 1000 frames with 25 Hz frame rate; stimulation data was obtained in 10 blocks à 100 frames with 12.5 Hz frame rate. **c.** Topview of a prepared mouse ready for imaging and display of one image frame. **d**. Exemplary vital parameters (heart rate (HR), respiratory rate (RR)) during data acquisition of one animal. **e.** Spectrogram of unfiltered raw signal of the same acquisition as in d. Mean HR and RR over time are indicated by white crosses **f.** Mean HR and RR during resting-state imaging in Hz; range of frequencies kept after filtering is indicated by black lines (n=6; >2recordings each). **g.** Exemplary signal time course of one pixel before and after preprocessing; black line on top of raw data shows the polynomial trend (order=5). Preprocessing included normalization and bandpass filtering from 0.1 to 1.0 Hz. **h.** Spatial registration process. Grey rectangles represent different data acquisitions of one mouse; An average across 8 acquisitions of one mouse is shown next to the schematic drawing, resulting in a blurred image due to misalignment. The first acquisition was repositioned with the bregma in the center and midsagittal line vertically oriented. Subsequent acquisitions of the same animal were aligned to the first acquisition by means of anatomical structures. The red line delineates the general mask. Red crosses depict references in virtual space. **i.** Schematic drawing of entire mouse brain, indicating position of bregma and general mask.

### Resting-state paradigm

Animals were anesthetized and fixed in a stereotactic frame to eliminate head motion. During the entire procedure the eyes were shielded from light. Images were recorded at a frame rate of 25Hz. During the anesthesia study we recorded 200 seconds, resulting in 5000 frames per acquisition, in both other studies we recorded 240 sec (=6000 frames) per acquisition (**Fig. 1b**). In a subset of mice (n=6) heart rate and respiratory rate was acquired simultaneously with calcium images, using the PowerLab data acquisition system (PowerLab 16/35, ADInstruments, New Zealand) and LabChart (LabChart, ADInstruments, New Zealand) (**Fig. 1d-f**).

### Preprocessing of imaging data

All data processing steps were performed in MATLAB (R2016b, The MathWorks, USA). Aims of our preprocessing steps were to reduce noise and signal fluctuations such as signal decay due to GFP bleaching by the light beam during acquisition and signal fluctuations induced by heart and respiration rate. Calcium images were first resized by the factor of 2/3, using the MATLAB function imresize, resulting in an image matrix of 220×220 pixels and resolution of 18.5 pixel/mm. Then resting-state data was intensitiy normalized. This was done using the mean fluorescence intensity of the signal time course (F) as reference intensity to compute fluorescence signal difference of every time point (ΔF) which was subsequently normalized (ΔF/F). Second, we applied a bandpass-filter with a passband from 0.1 to 1 Hz. This filter was implemented using the MATLAB functions cheby1 and filtfilt to design a Chebyshev Type I filter of order 2 and to perform zero-phase digital filtering. In a test data set heart and respiratory rate were always between the upper cutoff frequency of our filter (1 Hz) and the Nyquist frequency of 12.5 Hz, allowing their alias free removal by the bandpass filter (**Fig. 1d-f**). The lower bound of the filter was used to remove low frequency components such as the signal decay due to GFP bleaching. Filtered signal time courses were then cropped 10 seconds from beginning and end to reduce possibly remaining filter artifacts. An exemplary raw signal and corresponding preprocessed signal time course is shown in **Fig. 1g**. After preprocessing, videos were generated (**Suppl. Video 1**) to evaluate the recording and preprocessing quality. An experienced rater reviewed the video of each recording for motion artifacts, and excluded affected acquisitions. This resulted in 27 (out of 30, 10% excluded) acquisitions for 1.5% ISO, 30 (out of 41, 26,8% excluded) acquisitions for 1.0% ISO and 25 (out of 34, 26,5% excluded) acquisitions for MED+ISO protocol during anesthesia study. In the stroke study, the final number of acquisitions was 256 (out of 279, 8.2% excluded).

### Spatial registration

Preprocessed images from different days and animals were spatially registered to ensure intra- and interindividual correspondence of the analyzed cortical regions. To this end, all images were repositioned with their bregma in the center of the image matrix and the sagittal suture in a vertical line. This was achieved in one step for the first acquisition of every individual mouse by manually rotating and translating it in the image plane into a virtual space (3 degrees of freedom). For all following acquisitions of the same individual mouse, two steps were needed. In a first step images were manually aligned to the first (non-repositioned) acquisition by means of vasculature and anatomical landmarks. The hereby estimated transformation matrix was then combined in a second step with the transformation matrix associated with the spatial repositioning of the first acquisition, in order to reposition all acquisitions in the same way and—importantly—with only a single interpolation step (**Fig. 1h**).

### Masking of images

An individual mask was created for every acquisition in a two-step process. First, a general mask was used to limit the analysis to cortical areas in the focal plane and exclude more laterally located cortical areas, which were out of focus due to the curvature of the cortical surface. The masking area was chosen by comparing the registered images from different animals and covered a symmetric area in both hemispheres (**Fig. 1i**). Secondly, an individual mask was computed which excluded all pixels saturated due to autofluorescence, e.g. in areas affected by the infarct. For every acquisition the two masks—general mask and individual mask—were combined. These resulting masks were used for all following analyses to identify included pixels and their signal time courses. The saturated pixels were used for lesion size estimation and depiction.

### Power spectrum calculation

The periodogram function in MATLAB was utilized to calculate power spectral densities (or simply power spectrum). In order to construct spectrograms, the power spectral density was computed within a moving time window of 250 frames (=10sec), moving frame-by-frame over the entire signal time series. Spectrograms were calculated for the raw signal time courses of 6 randomly but consistent preselected pixels and subsequently averaged across these 6 pixels. An exemplary spectrogram and heart and respiratory rate for the same acquisition are depicted in **Fig. 1e**.

In the anesthesia study, the power spectrum was calculated for the preprocessed signal of three randomly but consistent preselected pixels in the right hemisphere and then spectral edge frequencies (SEF) and median frequency (MF) were derived as the frequencies at which the cumulative power reached respectively 5% (SEF 5), 50% (MF) and 95% (SEF 95) of the total power. A paired T test with subsequent Bonferroni correction was used for statistical testing of group differences.

### Approximate entropy estimation

In order to select an optimal anesthesia protocol in the anesthesia study and to ensure moderate levels of anesthesia with comparable neuronal activity throughout experiments, approximate entropy (ApEn) was estimated. ApEn was computed for the preprocessed signal time series of three pixels located in the right hemisphere, using the MATLAB file exchange function calculating fast approximate entropy, offered by Kijoon Lee (13). We decided to use the commonly used embedding dimension m=2 (14). ApEn was then calculated for a range of tolerance levels (tolerance=r*std, with the coefficient r being varied between 0.01 to 0.3 by steps of 0.01, and std being the standard deviation of the signal time series). The resulting ApEn scores were averaged across the three pixels, but separately for each tolerance level. Then the maximum approximate entropy (ApEn_max_) was determined among all tolerance levels to obtain an appropriate value for every data set as discussed by Chon et al (14). Paired T tests and subsequent Bonferroni correction was used for statistical testing of group differences in the different anesthesia protocols.

Based on comparisons between the different anesthesia protocols in the pilot experiment, we defined a threshold for ApEn_max_ equal to 1.25 and excluded all recordings from the stroke study with ApEn_max_ below this threshold (4 out of 279 acquisitions =1.4% of the recordings).

### Independent vector analysis

Independent vector analysis (IVA) as implemented in the Group ICA Of fMRI Toolbox (GIFT v3.0a; http://mialab.mrn.org/software/gift) was used to derive functionally independent cortical networks. IVA is an extension of ICA enabling its application to group data (15–17) and was recently suggested as an alternative to the widely used group independent component analysis (GICA), as it could be shown to better preserve subject variability (18). We used the algorithm to conduct both single subject analysis and cross-sectional group analysis in naïve mice. The number of components to be estimated by the algorithm has to be preselected. We used 20 components for single subject analysis and repeated the group analysis with increasing numbers of components (i.e. 16, 18, 20 and 30).

Single subject IVA was calculated for baseline data to locate the motor cortex for every mouse individually. The motor cortical region of the forepaw was identified by selecting the component, whose spatial map included a functional area located rostrolateral to the bregma. The coordinates of the peak pixel—the pixel with the highest correlation of its signal time course to the signal time course of the component—of this area were further used for motor cortex stroke induction by photothrombosis.

### Definition of functional networks and ROI using IVA

In order to find functional cortical areas across all mice, a meta-analysis of several group IVA analyses was conducted. To get the most robust results 16 group IVAs were calculated. In particular, we analyzed data from two different large groups of naïve animals (n=41 and n=47), and each animal was recorded on two independent days over 4 minutes (=6000 frames). Hence, we received four datasets by keeping the first and second recoding separate. For each of these four datasets, we conducted four different group IVA by altering the number of requested components (n=16, 18, 20, 30). We selected 10 out of these 16 analyses based on the criterion that IVA components should be consistently spatially represented across all group IVAs. In a second step, only individual components which were represented in at least 9 out of the 10 analyses were considered as robust, while less prevalent components were discarded for further analyses. The areas of the selected components were merged and the median of their peak pixel obtained. We used the median peak pixel in the sensorimotor cortex of both hemispheres to define circular regions of interest (ROI) centered on them, with a diameter of 8 pixels (≈0.43 mm) and containing a total of 49 pixels. These ROI were then used in further functional connectivity analyses. Also these ROI were masked, as described above. A ROI was only included in an analysis if more than 75% of its pixel were located within the mask. The ROI signal time courses were calculated as the average signal time course of all pixels within the corresponding ROI.

### Functional connectivity analysis

In order to evaluate the effect of anesthesia on functional connectivity in the anesthesia study and to characterize network changes in the brain after stroke in the stroke study, we conducted three types of connectivity analysis: seed-based analysis, ROI pair-wise analysis, and global connectivity analysis (only used in the stroke study).

For all three types of analysis, functional connectivity was computed as Pearson’s correlation coefficients between signal time series of different ROI (as resulting from the IVA meta-analysis) or single pixels. Correlation coefficients were further Fisher z-transformed, in order to allow parametric statistical testing and more accurate evaluation of high connectivity levels.

For seed-based functional connectivity analysis, connectivity scores were calculated between a selected ROI in the right caudal forelimb (rCFL; here considered the seed of analysis) and the signal time series of respectively all pixels on the cortical surface included within the mask. This analysis resulted in a map of connectivity scores. For ROI pair-wise analysis, connectivities were calculated between selected pairs of ROI and represented both as graph plots and as matrices. In graph plots, connections were represented as lines between ROI center-coordinates displayed topographically on top of an exemplary masked mouse cortex.

In the anesthesia study, connectivity scores were averaged across trials within condition, but separately for each individual mouse. Paired T tests and Bonferroni correction were used to compare these average scores between conditions. For visualization, these scores were further averaged across mice within each anesthesia condition. In the stroke study, two sample T test and Bonferroni correction was used for statistical comparison between stroke and sham groups at each time-point. If a ROI was located within or at the edge of a cortical area affected by autofluorescence, the connectivity scores calculated for this ROI were excluded from the analysis. If this masking procedure caused the sample size to drop for a specific ROI-pair or ROI-pixel-pair below 5 mice per group (i.e. stroke or sham group), this pair was excluded from analysis. For visualization, connectivity scores were averaged across mice within each treatment group.

Global connectivity analysis (19,20) was used to investigate overall functional connectivity alterations caused by the ischemic infarct in the stroke study. Global connectivity was computed for each pixel inside the contralateral hemisphere, as the average functional connectivity of this pixel with all other pixels inside the same cortical area. For statistical analysis of global connectivity alteration after infarct, baseline global connectivity scores were subtracted from the global connectivity scores at the days after stroke or sham procedure, for each mouse individually. For visualization, these differences in global connectivity were averaged within group and displayed in form of a topographical map for each time point. Instead, for statistical analysis these scores were further averaged across pixels, in order to receive one score per mouse and time point, and compared at each time point between groups using two-sided T tests with Bonferroni correction. To further explore global connectivity alterations, the histogram of the distribution of non-normalized global connectivity scores across pixels was obtained at Baseline, D1 and D28 after stroke. For this purpose, global connectivity scores were averaged across mice per time point and pixel, for the stroke and the sham group respectively.

### Stimulus dependent paradigm

Stimulation was performed subsequent to resting-state data acquisition at 12.5 Hz frame rate in a group of naïve animals (n=23). Acupuncture needles (Hwato, Ternimed, Germany) were placed subcutaneously between second and third digit of each paw. Stimulus trigger was given by the Vdaq software (Optical Imaging Ltd, Israel), passed on to a Train/Delay Generator (DG2A, Digitimer Ltd, UK). The final stimulus was generated by a constant current isolated stimulator (DS3, Digitimer Ltd, UK). Paws were stimulated alternating and each paw was stimulated for at least two imaging blocks. Each block contained of 10 independent trials. Every trial lasted 8 seconds and started with a 2 seconds baseline period followed by 4 seconds of stimulation with maximum 1.6 mA amplitude, 300 ms duration and 1 Hz interval, and concluded with 2 seconds of resting period (**Fig. 3 c**).

### Stimulation evoked activity

Signal time series recorded during paw stimulation were first normalized by subtracting and dividing through the average activity during the 2 second baseline period at trial start. The normalized time series were then averaged across trials, per each individual mouse and stimulated paw. In pilot studies, a stimulation-to-peak response delay measured approximately 380ms. Therefore, frames recorded within a time window of 300-460ms after stimulus onset (peak of response) and frames recorded within a time window 780-940ms after stimulus onset (trough of response) were averaged, and then the peak to trough amplitude calculated as the difference between these averages. The peak to trough amplitudes for stimulations at different days were averaged per condition and mouse. Subsequently the interindividual mean per condition was calculated. Then, all pixels with a mean peak to trough amplitude above the 98th percentile were determined and considered as the somatosensory core region representing the stimulated paw area in the somatosensory cortex.

### Photothrombosis

Photothrombosis was used to induce cortical focal ischemia. We performed a baseline *in vivo* calcium imaging resting-state acquisition at least one day prior to stroke induction and calculated a single-subject IVA to obtain the motor cortex coordinates for each individual mouse. Animals received 10 μl/g body weight of 1% Rose Bengal dye in saline intraperitoneally (i.p.) 5 minutes prior to anesthesia (isoflurane in 30%O2/70%N2O). The mouse was then placed into a stereotactic frame. A laser (Cobolt HS-03, Solna, Sweden) with a fiber optic bundle of 1.5 mm diameter at the tip was used to induce a lesion restricted to the area of interest and the rest of the cortex was shielded from laser light. The infarct was induced through the coverslip, dental cement and intact skull by illuminating for 17 minutes at constant 561nm wavelength and 25 mW output power at the fiber. Sham procedure was performed analogous, but without laser illumination.

Twenty-four hours after stroke induction, infarct area was measured via laser speckle contrast imaging using the PeriCam PSI System (PeriCam PSI System, Perimed, Sweden). Blood perfusion of the cortex was measured during 30 seconds and an averaged color coded picture obtained. The area with no and strongly reduced cortical blood perfusion was determined manually using ImageJ software (Version 1.49c, Fiji) (21).

### Neuroscore

The neuroscore was assessed as previously published (22). Briefly, the score is composed of the assessment of several subtests of both global and focal deficits. Assessment of global deficits included grooming, status of ears and eyes, posture, spontaneous activity and epileptic behavior. Focal deficits were evaluated by gait, grip, forelimb-asymmetry during tail suspension, circling behavior of both entire body or only forelimb, body symmetry and whisker response. Total score ranges from 0 to 54 points (26 point for general and 28 for focal deficits), higher score indicating worse deficits. Data was acquired once before stroke and on day 1, 3, 7, 14, 21 and 28 after photothrombosis. We used 2-way-ANOVA and Dunnett’s correction for multiple comparisons for statistical testing.

### Beamwalk test

The beamwalk test was performed by modifications as previously described (23). We used a 5×20 mm wooden beam. After training the mice at least 3 three days until successfully traversing the beam, a baseline run was recorded, after stroke data was acquired for day 3, 7, 14, 21 and 28 after stroke. On each testing day, every mouse had to run three times with a 30 second break in between. All trials were video recorded and analyzed frame by frame. Foot faults of the hindlimbs were counted for each paw separately and averaged for all three runs per time point. We used 2-way-ANOVA and Dunnett’s correction for multiple comparisons for statistical testing.

### Statistical analysis

Statistics for behavior analysis have been calculated using Graph Pad Prism. Statistics for *in vivo* calcium imaging experiments have been calculated in MATLAB. The applied statistical test is specified in the respective Methods section.

### Data

Data will be made available on reasonable request to the corresponding authors.

## Results

The overall aim of this study was to establish a translational tool adopting analysis paradigms common in fMRI connectivity studies to an optogenetic neuronal activity imaging approach in mice. To achieve this, we performed a stepwise study design in three sub-studies, first analyzing a stringent mouse sedation protocol, then establishing network analyses with this novel tool and finally testing in an experimental stroke model. The anesthesia study aimed to establish a sedation protocol allowing for stable and reproducible recordings of resting-state neuronal activity and to derive physiologically meaningful functional connectivity scores. The network study aimed to define functional cortical networks and to identify nodes in these networks which can be used as ROI for functional connectivity analysis in mice. The stroke study aimed to assess functional connectivity alterations after experimental stroke using the paradigm developed in the anesthesia and network studies.

### Anesthesia study: Determining the impact of anesthesia on functional connectivity analysis

We tested 3 protocols (n=9; ≥3 acquisitions per protocol) differing in the depth of anesthesia: deep anesthesia with 1.5% isoflurane inhalation (1.5% ISO), low anesthesia at 1.0% isoflurane inhalation (1.0% ISO) and light sedation with a combination of medetomidine injection and 0.75% isoflurane inhalation (MED+ISO). The impact of anesthesia was already apparent in the preprocessed signal time courses and computed power spectra for the three anesthesia conditions (**Fig. 2a, b**). Moreover, both the spectral edge frequency (95th percentile, SEF95) and the maximum approximate entropy— a measure of randomness—differed significantly between all three anesthesia groups (**Fig. 2c, d**). The lowest entropy was observed in mice receiving the deeper anesthesia (1.5% ISO). We also investigated directly the impact of the different anesthesia protocols on functional connectivity. In a seed-based analysis, connectivity was calculated between a ROI of the sensorimotor cortex and the rest of the cortex (**Fig. 2e**). In this analysis considerable differences in functional connectivity were apparent. As the combination of ISO+MED preserved the largest degree of data entropy, largest spread in the frequency spectrum and was subjectively the most stable and reliable to handle protocol, this approach was used for the subsequent sub-studies: the “network study” and the “stroke study”.

**Figure 2:**
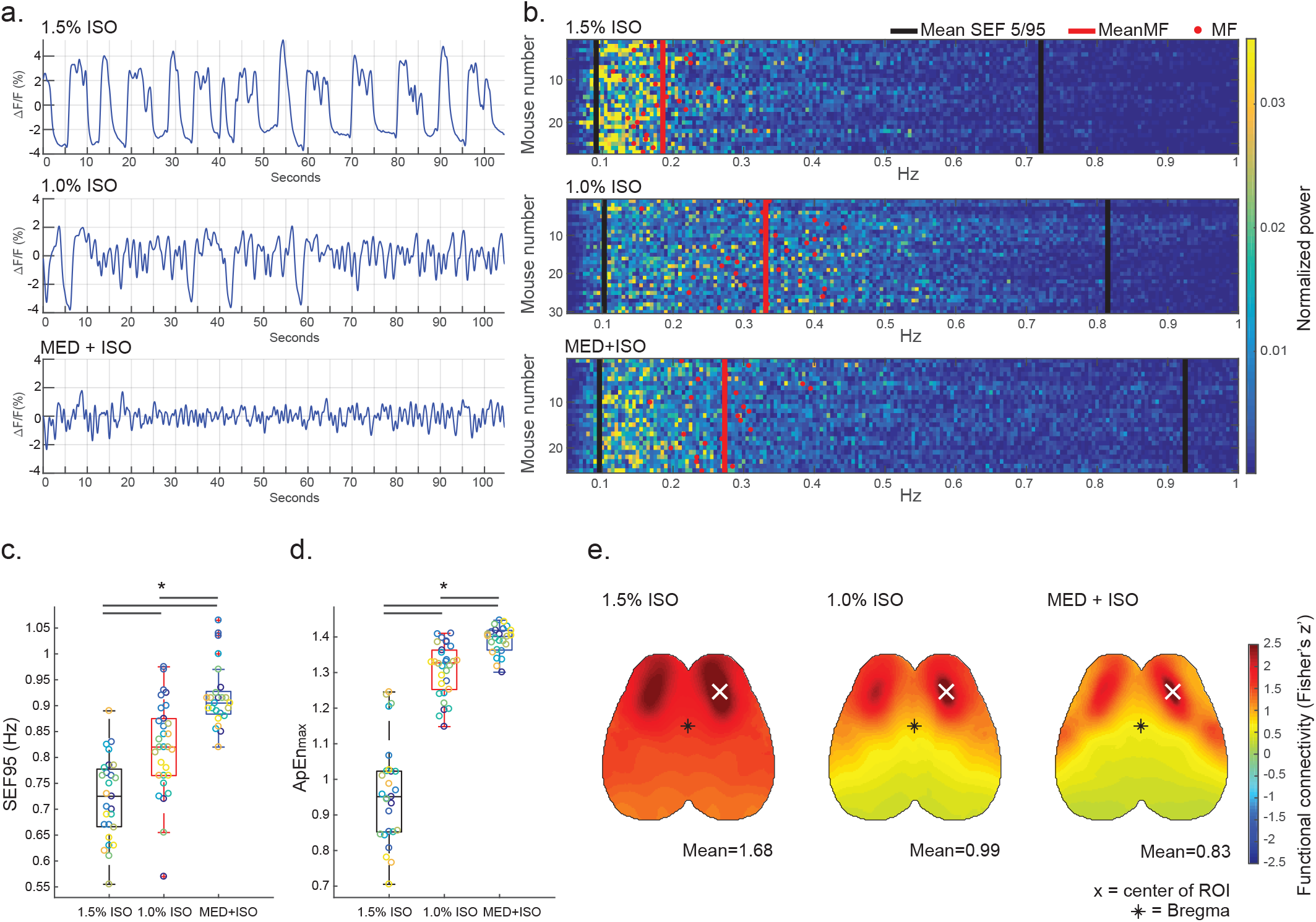
Influence of anesthesia depth on calcium recordings and functional connectivity. **a.** Representative plots of filtered signal time course for all three anesthesia conditions. **b.** Power spectra for all trials in each anesthesia condition, normalized to the total power. Lines display median (red) and spectral edge frequencies (black) of the 5th (SEF 5) and 95th (SEF 95) percentile. **c.** Mean SEF 95 and **d.** mean maximum approximate entropy (ApEn_max_) for each trial in the three anesthesia conditions. Each individual mouse is displayed in a different color. Significance bars above the plots indicate differences between conditions at a significance level α= 0.05. **e.** Topographic depiction of seed-based functional connectivity averaged across mice; *= Bregma, white cross indicates seed.

### Network study: Identification of functional cortical networks by independent component analysis

Our primary aim was to map functionally relevant cortical areas in the mouse brain to obtain coordinates to use for further seed-based and ROI pair-wise connectivity analyses. To retrieve cortical networks, we performed a meta-analysis of 10 group-IVA which originated from different data sets and parameters (for more details see method section). This meta-analysis resulted in 12 components, with matching spatial patterns in at least 9 of the 10 IVAs (**Fig. 3a, and Suppl. Fig. 1**). For each of these 12 components, the corresponding spatial maps from the different IVAs were overlaid and pixels being in at least four analyses over-threshold in all of these maps were marked as belonging to the cortical network described by that component (**Fig. 3b,** orange pixel). Furthermore, for each of these 12 components, the peak pixels in both hemispheres were determined in the spatial component maps from the different IVA and the median peak coordinates were subsequently calculated across the different IVA. The individual peak coordinates differed from the median peak coordinates on average by only 3.2 pixels (≈0.17 mm) (**Suppl. Fig. 1i**). The median peak coordinates were then used as center coordinates for ROI definition and interpreted as the center of the cortical areas involved in the functional networks captured by the IVA components (**Fig. 3b**). The function of these cortical areas was inferred by comparison of median peak coordinates to the Paxinos Brain Atlas (24)(**Suppl. Fig. 1 and Suppl. Table 1**). We further aimed to validate the biological significance of identified components performing a somatosensory stimulation of all 4 paws (n=23; **Fig. 3c**). We detected a specific response of the components identified as the sensory cortex (using the Paxinos atlas (24)) to the stimulus, whereas the components identified as motor cortex did not react to stimulation (**Fig. 3d**). Stimulation evoked contralateral activation of defined cortical areas, which were symmetrical when comparing left and right stimulation (**Fig. 3e**). An overlay of the components identified by IVA in the somatosensory cortex onto the areas of maximal activation in response to electrical paw stimulation (ΔF/F 98th percentile) revealed a high overlap (**Fig. 3f**). These findings indicate that the IVA analysis pipeline is able to perform unbiased identification of anatomically and functionally distinct cortical areas with neurophysiological function.

**Figure 3:**
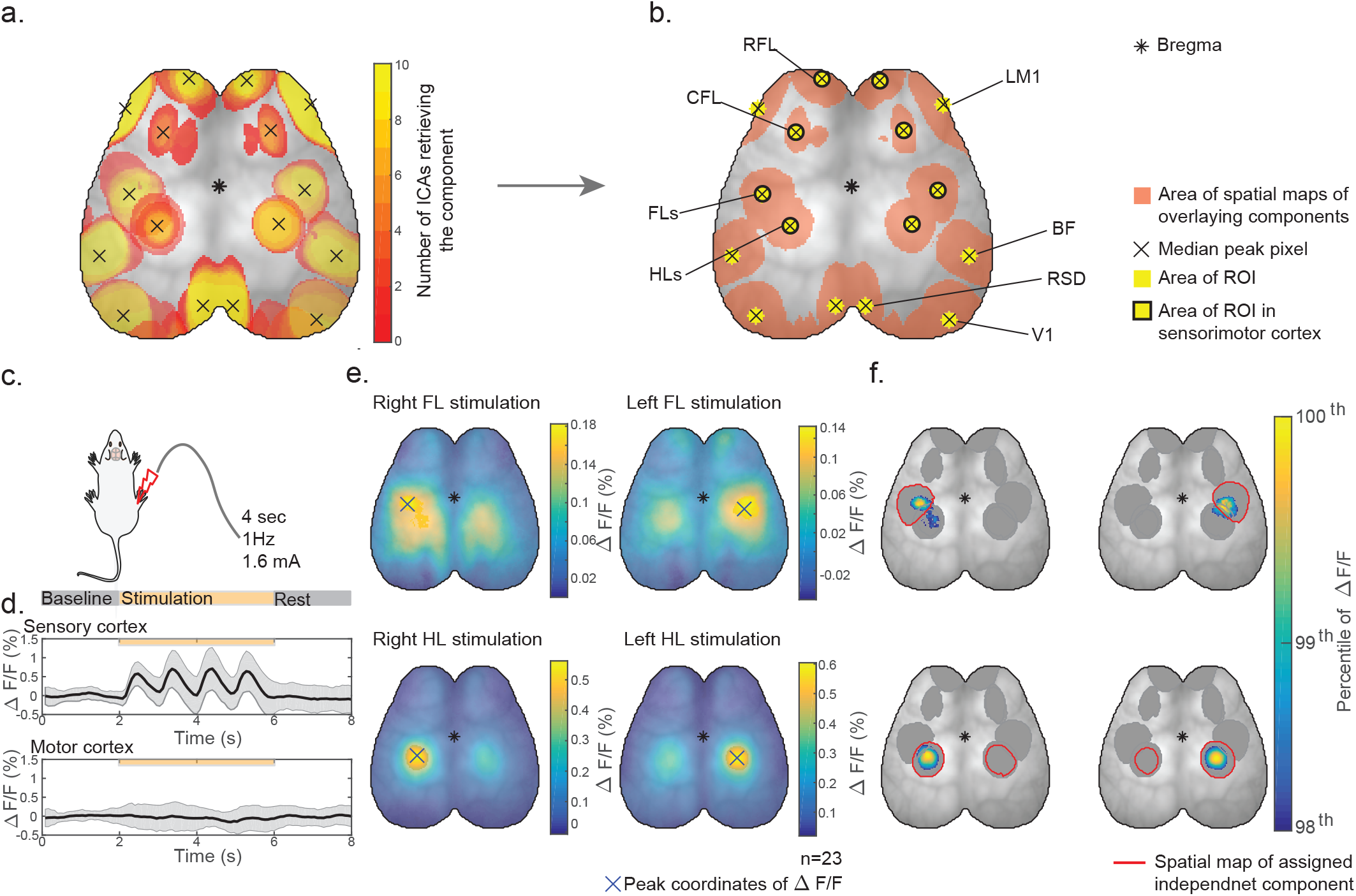
IVA derived functional networks and validation by somatosensory stimulation. **a.** Topographical heatmap of the 12 components retrieved by the meta-analysis of 10 group IVA. Pixel-color indicates, in how many of the group-IVA a given pixel was present. **b.** Topographical depiction of all regions of interest (ROI) (yellow). The centers of the circular ROI (crosses) were determined as the median peak pixels of the matching IVA components. ROI were anatomically assigned as rostral forelimb area (RFL), lateral M1 motor cortex (LM1), caudal forelimb area (CFL), forelimb sensory (FLs), hindlimb sensory (HLs), barrel field (BF), retrosplenium (RSD) and primary visual cortex (V1). **c.** Experimental setup of paw stimulation paradigm; 10 trials were recorded successively, one trial consisted of 2 seconds baseline activity, followed by 4 seconds of 1 Hz stimulation (with stimuli of 1.6mA and 300ms duration) and 2 seconds resting period. **d.** For the example of hind limb stimulation in one animal, the evoked response (averaged across trials) in the contralateral sensory (ROI in HLs) and motor cortex (ROI in CFL) is shown at the bottom of the panel; black line corresponds to the mean, grey area to the range of standard deviation. **e.** Topographical depiction of mean ΔF/F of 23 mice for each stimulated paw (right and left forelimb (FL) and right and left hindlimb (HL)) **f.** Overlap of the 98th percentile of mean stimulation data (color coded), the areas retrieved by IVA analysis (grey) and assigned component (red). In b, c, f and g the star indicates the bregma as a spatial reference.

### Stroke study: Assessment of functional connectivity after experimental stroke

As a last step, we applied our novel approach to assess pathological changes and monitor recovery of functional connectivity after an acute injury to the motor cortex. We induced a photothrombotic lesion in the motor cortex (n=23) which was identified by ICA of all individual animals. Control animals received sham surgery (n=17). We then performed widefield calcium imaging as well as behavioral testing on days 1, 3, 7, 14, 21, 28 and 56 after surgery. We could show a high correlation (R^2^=0.704; p<0.0001; n=23) of the area of saturated pixels and area of blood flow reduction assessed by laser speckle imaging on D1 after stroke (**Suppl. Fig. 2**). Hence, cortical autofluorescence of necrotic brain tissue was used to determine the lesion size and location as well as their temporal evolution until day 56 after stroke (**Fig. 4a**). Additionally we assessed deficits in commonly used behavior tests for post-stroke motor deficits in the photothrombotic lesion model (**Fig. 4b, c**). Deficits in motor function were clearly detectable during the acute phase (days 1-7 after stroke) in the composite neuroscore as well as the beamwalk test. However, while the neuroscore still identified a significant behavioral deficit up to 56 days, no significant deficits were detectable in the beamwalk test after 14 days post-lesion. Hence, behavior tests might be suitable to determine motor deficits in the acute phase after stroke but are less sensitive to analyze long-term recovery, stressing once more the need for *in vivo* imaging application such as our tool to measure recovery by more sensitive means.

**Figure 4:**
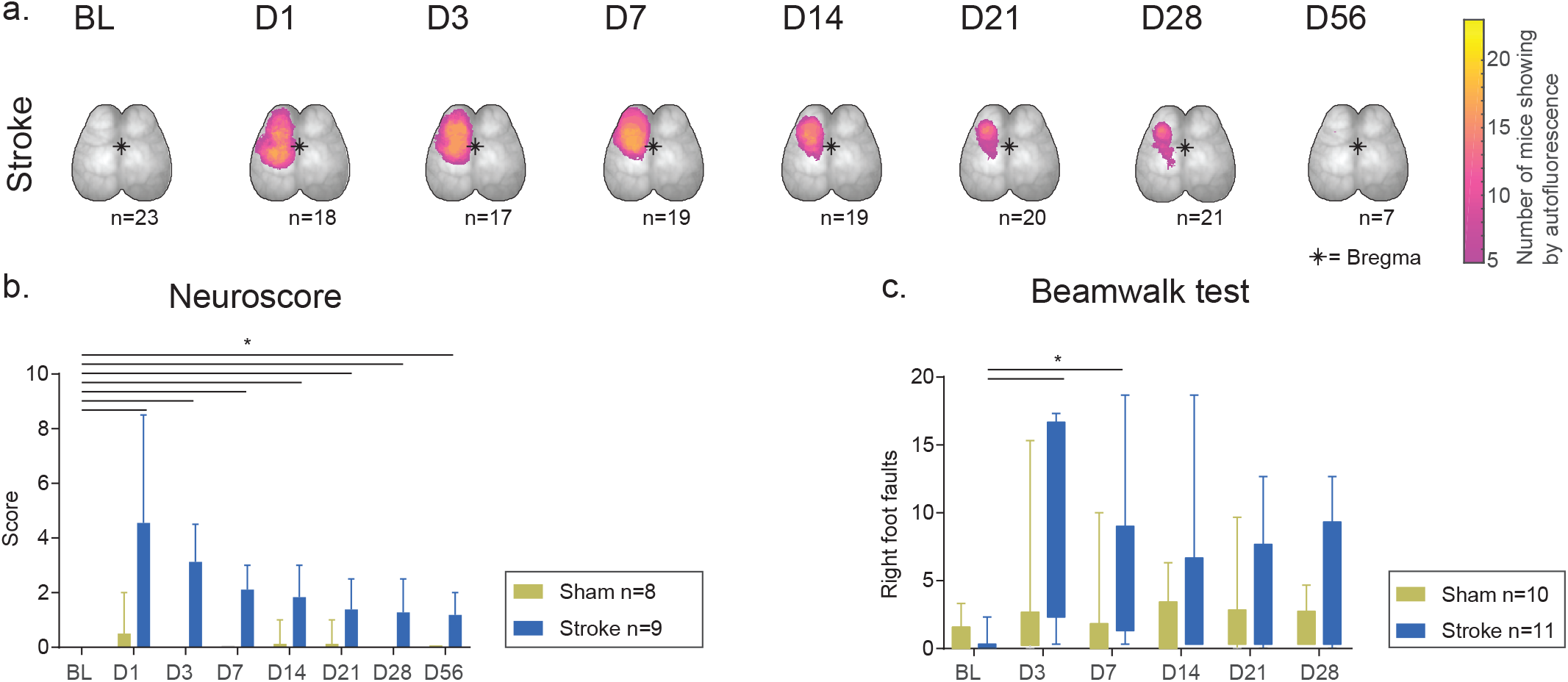
Lesion incidence maps and behavioral deficit after stroke. **a.** Lesion distribution based on cortex autofluorescence after stroke for each acquisition time point. **b.** Time course of the multi-parameter neuroscore (sham: n=8, stroke: n=9) and **c.** right foot faults counted during beamwalk test (sham: n=10, stroke: n=11). Statistical values have been calculated using 2way-ANOVA, *= p<0.05.

Analysis of functional connectivity was performed using the resting-state widefield calcium imaging paradigm, as established in the anesthesia and network sub-studies. Changes of functional connectivity scores over time allowed us to track pathological changes and restoration of network integrity after photothrombotic motor cortex lesions. In line with previous findings (25), we observed interhemispheric homotypic correlations as the dominant connectivity patterns in naïve brains (compare to Fig. 3 and BL images **Fig. 5a-c**). Therefore, we utilized the ROI corresponding to the structurally intact motor area contralateral to the lesion (the right caudal forelimb area; rCFL in Fig. 3b) as a seed to compute functional connectivity to all other pixels in the cortex (**Fig. 5a**). After infarction, we detected reduced interhemispheric connectivity of this rCFL ROI to the hemisphere ipsilateral to the infarct, in contrast to its rather unaffected intrahemispheric functional connectivity. The reduced interhemispheric connectivity persisted until 56 days post lesion. Additionally to this seed-based connectivity analysis, we also conducted ROI pair-wise analysis using the 8 ROI (4 in each hemisphere), determined by the group-IVA in naïve animals and being located in the sensorimotor cortex (see Results, second section, and Fig. 3b). Corresponding to the seed-based connectivity analysis, also this pair-wise analysis of ROI in functionally defined cortical areas revealed a loss in connectivity to the ischemic area resulting in broad network disturbances after infarction (**Fig. 5b-d**). Interestingly, homotypic connectivity of the structurally unaffected motor cortex frontal to the lesion area (RFL=rostral forelimb motor cortex) was diminished until 28 days after stroke. In contrast, connectivity to the somatosensory cortex caudal to the infarcted cortex area remained mainly unaffected after stroke despite a similar distance to the lesion core (**Fig. 5e, f**), indicating more pronounced disturbances of connectivity in functionally linked networks such as the motor cortex. Consequently, a complex pattern of connectivity time courses can be observed after stroke with partial reconstitution of connectivity to some areas, unaffected connectivities to more remote network components as well as sustained reductions in connectivities to the primarily injured cortex area (i.e. lCFL) (**Fig. 5e,f**). We additionally aimed to study more global consequences of acute injuries on cortical network function. We computed global connectivity to examine the overall functional connectivity of each pixel of the cortex relative to all other pixels (19,20). In this analysis, we reduced the masked area to include only the hemisphere contralateral to the infarct in order to examine the secondary impairment or compensatory changes in the non-lesioned hemisphere. We observed a significant transient increase in contralateral global connectivity at day 1 post injury which returned to comparable levels of the sham-operated control group (**Fig. 6a** and **Suppl. Fig. 3**).

**Figure 5:**
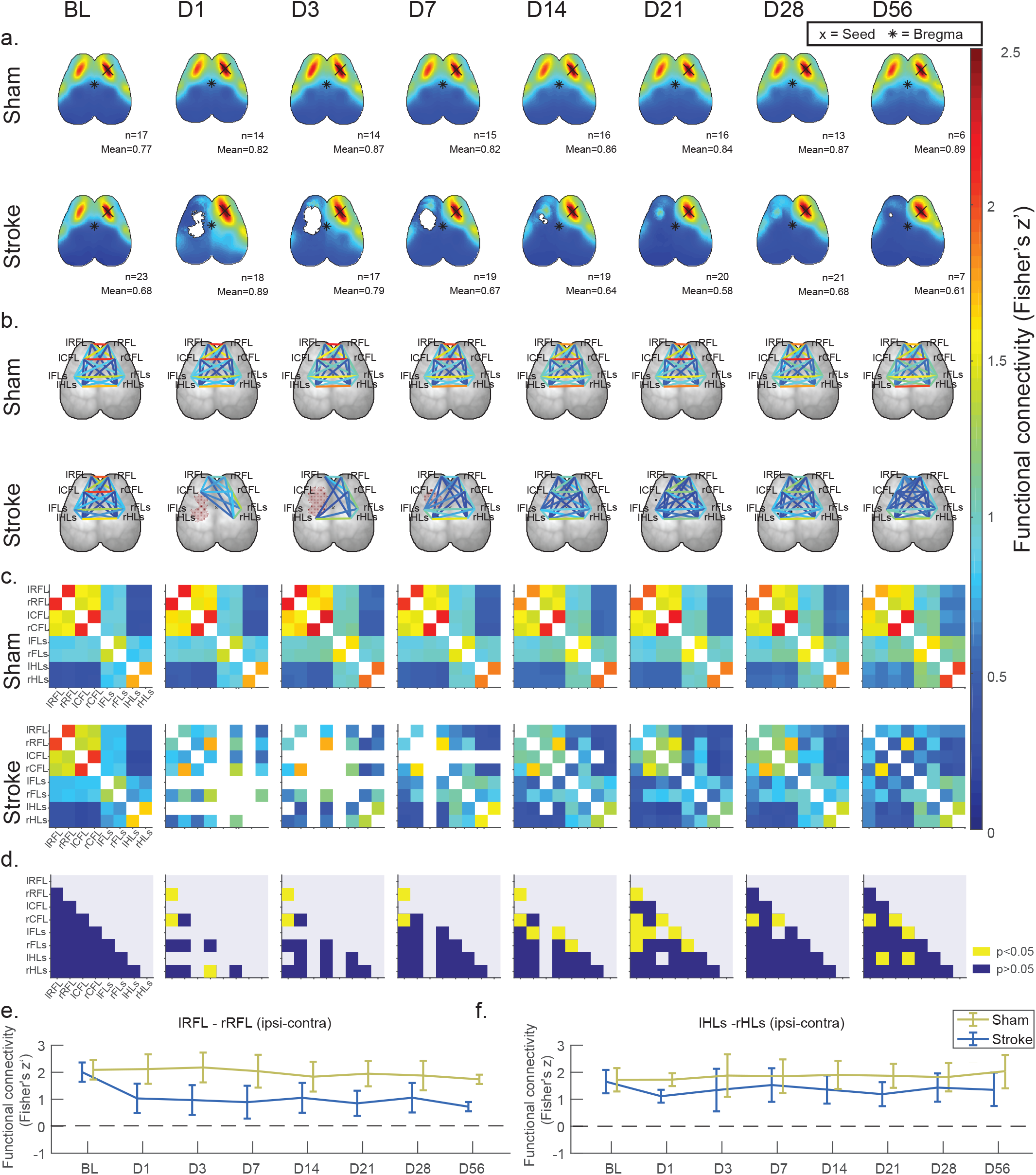
Changes in functional connectivity after stroke. Each column in panels **a.** to **c.** depicts functional connectivity scores (averaged within sham and stroke group) for respectively one day (BL: baseline; D1-D56: days 1-56 after surgery). **a.** Topographical depiction of seed-based functional connectivity. The seed pixel (X) is located in the right caudal forelimb (rCFL) cortex, the area contralateral to the infarct. **b**. Graph-representation and **c.** matrix-representation of functional connectivity between 8 ROI, 4 in each hemisphere placed in rostral forelimb (RFL), caudal forelimb (CFL), hindlimb sensory (HLs) and forelimb sensory (FLs) areas, r and l refer to right or left hemisphere respectively. Functional connectivity is displayed as Fisher z-transformed Pearson’s correlation; squares which contain less than 5 data points are not displayed. **d.** Results of T test between sham and stroke for each acquisition time point; p values were corrected via Bonferroni correction for multiple testing. Color codes indicate a significant group difference (p-corrected<0.05) in yellow. Functional connectivity between **e.** right and left CFL, and **f.** right and left HLs in sham and stroke operated groups. Data points and error bars represent mean and standard deviation.

**Figure 6:**
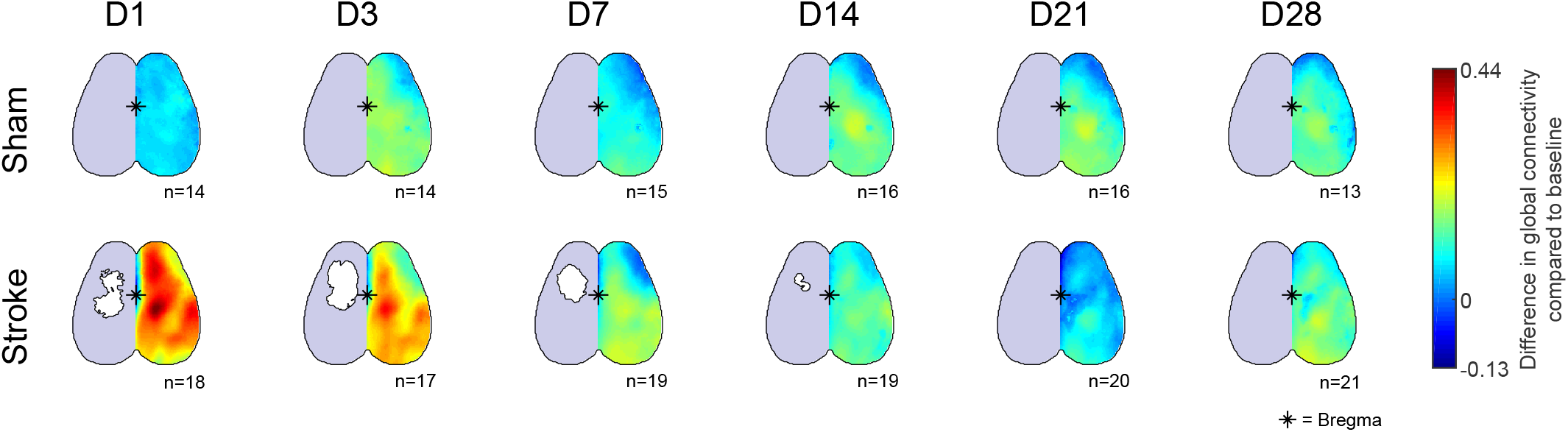
Analysis of contralateral global connectivity after stroke. **a.** Depiction of the mean change in global connectivity after surgery (D1-D28: days 1-28 after surgery) at each pixel in the contralateral hemisphere. Change in global connectivity was calculated by subtracting for each pixel the baseline global connectivity at the same pixel.

## Discussion

Taken together, our results demonstrate that our novel tool is a robust and highly sensitive method to determine changes in cortical functional connectivity after stroke. The developed anesthesia protocol and analysis algorithms allow the analysis of complex changes in functional connectivity during the time course of post-stroke recovery and provide a meaningful complementation to the conventional method to determine recovery in experimental stroke research by behavior testing.

Investigation of brain function by electroencephalography (EEG) and fMRI plays a key role in translational neuroscience and clinical neurology. These tools have increased our knowledge on healthy and pathological brain function in patients. However, comparable approaches to study function of the entire forebrain in laboratory mice are currently very limited. We combined and advanced the method of *in vivo* calcium imaging (3,4) with commonly used algorithms from fMRI and EEG analysis to fill in this gap of translational tools for characterization of functional networks. Our tool provides better spatial and temporal resolution of brain activity than fMRI in mice, at the expense of acquiring only cortical information. Additionally, this imaging modality allows high reproducibility, low stress levels for the mice by having short procedure times, and low cost compared to fMRI. We offer three major enhancements: An advanced analysis approach in conjunction with establishing of a rigorous protocol of minimal sedation for reproducible imaging conditions and spatial alignment of different recordings of the same mouse as well as recordings from different mice. Due to these advances our novel tool represents a substantial improvement over previously developed, similar approaches for *in vivo* widefield neuronal calcium imaging (3,4).

In order to successfully bridge preclinical models and clinical trials, a maximized comparability of research approaches, interventions and diagnostic tools is necessary. To overcome existing limitations in translational research and identify the key obstacles, several international research consortia have been established. Interestingly, most of these research consortia have so far focused on the methodological drawbacks in the design and reporting of preclinical research and proposed novel guidelines and study designs to improve this situation (26–28). However, ensuring that a biological effect does not become “lost in translation” relates not only to the way drugs are applied, animals are handled or data is reported. Comparability of the readouts is an equally critical measure that is currently still widely neglected (28–30). Therefore, our advanced tool addresses a crucial aspect by matching mouse optogenetic imaging to human fMRI by the use of comparable analysis algorithms which will allow a previously unprecedented comparability in the interpretation of functional neural networks between mice and men.

*In vivo* widefield calcium imaging is still a new method in neuroscience. In the past, various other functional neuroimaging modalities have been established for rodent research such as two-photon imaging, intrinsic optical imaging, fMRI and positron-emission tomography (PET). Two photon imaging provides very high resolution imaging of cortical structures including neuronal calcium concentrations, but are limited to only small cortical areas with up to maximally a few hundred neurons and additionally requires invasive craniotomy and implantation of cranial windows. In contrast, our tool offers acquisition of whole forebrain neural activity at the expense of losing single neuron resolution. This holistic approach on network function by our widefield imaging tool allows a more differentiated analysis of various functional domains of the cortical network in contrast to the widely used multiphoton imaging of small cortical areas. Additionally, our tool does not require trepanation of the skull and window implantation which might affect cortical function. Other often used imaging modalities such as positron-emission tomography (PET) and fMRI are still very challenging in small rodents due to the complexity and costs of such a setup (1,2). Although these modalities provide the possibility of investigating the entire brain and not only cortical structures, they lack both high spatial and temporal resolution. Moreover, fMRI and PET usually require deeper anesthesia compared to our tool, which will affect neuronal function as clearly demonstrated in our study. Thereby, widefield calcium imaging expands existing options such as two-photon imaging to display larger structures up to the entire forebrain cortex and has the potential to become a routine technique for studying network function in mice (3,4).

In the healthy brain localized neural activity and blood flow regulation are closely linked by a mechanism known as neurovascular coupling, which is a multicellular process encompassing neurons, glia and vascular cells (6,31). However, brain disorders such as Alzheimer’s disease and stroke impair physiological neurovascular coupling (6,32). This impairment plays a pivotal role in blood-flow-dependent imaging modalities such as fMRI or intrinsic optical imaging, because the possibility to draw conclusions on neural activity is based on the assumption of an intact neurovascular coupling. In contrast, our tool allows the blood-flow independent measurement of neural activity by directly measuring the signal (GCaMP fluorescence) from excitatory neurons.

An important consideration for the design of our imaging approach was to generate a widely applicable and relatively easy to use tool. Therefore, we chose to perform the imaging in anesthetized mice, even though awake-imaging is possible at expense of time investment to habituate the animals to experimental set-up (4). Anesthesia depth can affect neuronal activity, with deeper levels of anesthesia leading to slower activity with higher amplitude (33). This effect is likely to interfere with analysis of functional imaging data. During the process of finding an appropriate sedation protocol, we experienced imaging parameters to be very susceptible to the anesthesia depth. In contrast to some previous studies which reported comparable results in widefield calcium imaging between awake and anesthetized animals (4,34), our study revealed a profound impact of the anesthesia protocol on functional connectivity measures. We used the approximate entropy (ApEn_max_)—a parameter which was highly influenced by the tested anesthesia protocols and previously reported to be affected by anesthesia also in EEG recordings (35)—as a measure for selection of the anesthesia procedure with the least impact on physiological randomness of the signal time series. Thereby we identified light sedation to only minimally affect randomness as well as preserved a high spectral edge frequency the signal time courses which are well-established measures of depth of anesthesia (36). These are important findings with potential impact also on related imaging fields using anesthetized/sedated rodents, because the effect of anesthesia is widely disputed and discrepantly reported in the literature. While some studies reported—consistent with our findings—a substantial influence of anesthesia on functional connectivity (37–40), others did not observe such an influence in their datasets (4,34). This discrepancy might be due to differences in imaging modality, analysis strategy but also the sensitivity to anesthesia-dependent effects in the different approaches. Yet, the effect was unequivocally evident in our standardized and randomized analysis and needs careful consideration in future studies. Therefore, we established a highly reproducible protocol for the standardized light sedation of animals with medetomidine and low-dose isoflurane and provide here a very detailed and easily reproducible protocol for this light sedation approach.

ICA is a well-established method used with fMRI data to identify resting-state networks in the human brain (41,42). To define functional networks we applied ICA to our resting-state dataset, more specifically a multivariate extension termed IVA. IVA was introduced by Lee et al. and is an alternate way of group fMRI analysis which avoids permutation ambiguity of ICA (15). Identification of networks via ICA is independent of a priori seed definition, region of interest or model time course (18). In human fMRI and EEG, it is well established that components retrieved by ICA are mostly overlapping with brain areas activated during tasks (43). To our knowledge we are first to validate this paradigm also in a rodent imaging approach by demonstrating overlap of cortical areas identified by ICA and somatosensory stimulation. Similar to the experiments in humans, spatially overlapping results from stimulation and ICA analysis confirm the involvement of areas detected by ICA in particular sensory or motor functions. In comparison, existing imaging methods usually derive coordinates of ROI from sensory evoked responses or the stereotactic coordinates retrieved from brain atlases (3, 44–47). In contrast, ICA-derived coordinates provide functional ROI, where ROI determination and experimental investigation of associated parameters such as functional connectivity can be accomplished in the same *in vivo* experiment.

As a proof-of-concept application of our tool in a disease model, we chose to investigate brain network changes after cortical ischemia in the motor cortex. We detected a transient increase in contralateral global connectivity in stroke mice. This observation is in accordance with a previous study performed in rats using fMRI (46) and was also reported in human stroke patients (48,49). As previously reported we found very strong functional connectivity between homotypic areas of both brain hemispheres under healthy conditions (25), which was significantly reduced after stroke in the acute and subacute phase. In our model the observed changes in functional connectivity remained localized to areas of the motor cortex but did not affect the somatosensory cortex. This post-stroke desynchronization in neural activity might mirror changes due to both the acute injury and starting regenerative processes as well as secondary degeneration due to acute neuronal damage (50). Interestingly, a similar pattern of confined changes to closely associated cortex regions was previously also observed using stimulus-dependent imaging after cortical injuries (51) while subcortical lesions seem to have a more widespread impact also on distant functional systems such as both sensorimotor and visual system after subcortical stroke (46). At later time points we see improvement of interhemispheric functional connectivity which might be expression of restoration of bilateral brain communication and therefore a marker of regeneration. With regeneration of brain functional connectivity we see amelioration of behavioral deficits.

Rodent behavior testing after acute brain injuries, however, has numerous limitations and is— although widely used in the research community—a rather insensitive and unspecific readout for neurological recovery (52–54). Besides the intrinsic limitations to assess complex neurological deficits by simplified tests in small rodents, also the interpretation of behavioral test results is largely rater-dependent and biased by the selection of the used tests. On the contrary, *in vivo* imaging of functional cortical networks provides unbiased, sensitive assessment of the recovery process. Moreover, our tool offers the possibility to analyze the nearly the entire forebrain cortex and hence different functional cortical systems simultaneously. Findings such as the involvement of different cortical functional systems, their degree of impact on regenerative processes or possible transient compensatory mechanisms, can then direct more in-depth analyses of the identified cortical systems of interest.

With *in vivo* widefield calcium imaging in combination with advanced analytical strategies such as ICA, we present an easy to handle tool for repetitive study of functional cortical networks in mice. Particularly the comparability of analysis algorithms with human fMRI approaches will allow now urgently needed translational imaging studies from mouse to men for mechanistic studies and development of novel therapies.

## Acknowledgements

We thank Prof. Timothy Murphy (University of British Columbia, Vancouver, Canada) for constructive discussions on this tool. This work was supported by the Excellence cluster of the German research foundation “Munich Cluster for Systems Neurology (SyNergy)” and the German Research foundation (DFG, LI-2534/2-1 to A.L. & DU 1626/1-1 to M.D.)

